# Single-Cell RNA Sequencing and Inferred Protein Activity Analysis Reveal a Distinct Tumor Phenotype in Early-Onset Colorectal Cancer Patients

**DOI:** 10.1101/2025.05.15.653154

**Authors:** David A. Cohen, Aleksandar Obradovic, Luka Jovanovic, Yosuke Mitani, Tianxia Li, Soobeom Lee, Dechokyab De, Casidhe-Nicole Bethancourt, Alice Shin, Karen Dunbar, Francesco Cambuli, Christopher Lengner, Ning Li, Caitlin Rogers, Jennifer Ho, Viktoria Jakubikova, Emma Blystone, Yoanna Pumpalova, Beatrice Dionigi, Sanja Vickovic, Chia-Wei Cheng, Joel Gabre

## Abstract

Colorectal cancer (CRC) diagnosed before age 50 years (early-onset CRC, EO-CRC) is rising at an alarming rate, yet its molecular and microenvironmental drivers remain poorly understood. EO-CRC is highly heterogeneous, and while subtle differences from late-onset CRC (LO-CRC) have been reported, their full extent remains unresolved due to the limited scope of previous studies. Here, we integrate public data with in-house clinical samples profiled by single-cell RNA sequencing (scRNA-seq) and multiplex immunofluorescence (mIF) to compare EO-CRC and LO-CRC. Additionally, we employ gene regulatory network-based protein activity inference (VIPER), enabling a more precise characterization of key regulatory proteins driving tumor-stroma interactions. Our analysis reveals that EO-CRC and LO-CRC have a largely similar immune composition, challenging previous reports of an “immune-cold” phenotype in EO-CRC. However, we identify distinct stromal differences, including a significant enrichment of fibroblasts in EO-CRC. Notably, we define a previously unrecognized epithelial subpopulation in EO-CRC, marked by high expression of toll-like receptor 4 (TLR4) and C-C chemokine receptor type 5 (CCR5)—key mediators of inflammation-driven tumor progression and fibroblast recruitment. These findings suggest that EO-CRC may be driven by a tumor-intrinsic inflammatory phenotype with enhanced stromagenesis, providing new insights into potential mechanisms underlying its increasing incidence in young adults.

## Introduction

Colorectal cancer (CRC) is a major health burden, ranking as the second most common cancer and the third leading cause of cancer-related death in the United States (US) among individuals under 50 years old (early-onset CRC, EO-CRC). While CRC incidence and mortality have decreased in those 50 and older (late-onset CRC, LO-CRC) during the past 30 years, the opposite trend has been observed in younger adults. In the US, EO-CRC incidence rose from 8.6 to 12.9 per 100,000 people between 1992 and 2018, and a similar upward trend has been observed in Europe ^1, 2^. This demographic shift is even more pronounced in younger age groups, with the most significant increases seen in those aged 20-39 ^2^. Notably, left sided colon and rectal cancers have been the primary driver of this rise in EO-CRC ^3^. Current projections suggest that within the next decade, EO-CRC may account for 25% of rectal cancer cases and up to 12% of colon cancer cases in the US ^4–6^.

While certain well-known risk factors, such as a family history of CRC and inherited syndromes (e.g., Lynch syndrome and Familial Adenomatous Polyposis, FAP), play a role in EO-CRC, these account for only 25-30% of cases ^7, 8^. The majority of EO-CRC cases are sporadic, with no clear familial or genetic predisposition. This highlights the need for a deeper investigation into other potential contributing factors, both genetic and environmental. The birth-cohort effect has been used to describe the rise in sporadic EO-CRC, as data suggest individuals born after the 1950s exhibit a higher age-adjusted risk of CRC than those born in earlier generations ^2, 9–11^. Rising rates of obesity, dietary changes, and lifestyle factors, such as physical inactivity and alcohol consumption, have been implicated as possible etiologies for this birth-cohort effect ^12^.

At the molecular level, sporadic EO-CRC represents a genetically and pathologically heterogeneous group of cancers, similar to LO-CRC ^13, 14^. Recent investigations into the tumor microenvironment (TME) have revealed subtle differences with several studies suggesting that EO-CRC tumors may exhibit an immune “cold” phenotype, characterized by reduced T cell infiltration and diminished immune surveillance ^15, 16^. Despite these emerging insights, a comprehensive biologic signature of EO-CRC remains elusive, and the mechanistic links between suspected environmental risk factors and tumor development have yet to be established.

To date, only one single-cell RNA sequencing (ScRNA-seq) study has compared the molecular landscape of EO-CRC to LO-CRC, analyzing 14 patients (7 EO-CRC, 7 LO-CRC) from an East Asian cohort, which may limit broader applicability ^16^. This study is further limited by a high rate of data dropout, common to ScRNA-seq data. To address these limitations and provide deeper molecular insights, we present a comprehensive single-cell analysis of EO-CRC, examining 42,931 cells from 30 patients (11 EO-CRC vs 19 LO-CRC) from a Western cohort and utilize a novel gene regulatory network-based algorithm for inference of transcription factor and signaling protein activity from expression of their downstream targets (VIPER). This algorithm has previously demonstrated improvement in clarity of gene expression for analysis of ScRNA-seq data by increasing cluster resolution and amplifying biological signal-to-noise for activity of key signaling proteins and regulators of cell state ^17, 18^. Indeed, this protein activity inference approach previously applied to bulk RNA-sequencing data has led to a Clinical Laboratory Improvement Amendments (CLIA)-approved clinical test matching aberrant protein activity in tumors to druggable proteins ^19^. When applied to single-cell data, this protein activity inference approach leads to dramatic improvement in signal-to-noise by inferring activity of upstream signaling proteins and transcription factors from weighted expression of >100 downstream target genes. Even with high rates of gene expression dropout inherent to ScRNA-seq, enough downstream genes are typically retained to infer upstream regulatory protein activity, thereby transforming a typical ScRNA-seq gene expression matrix with 20,000 genes and 80% missing data to an inferred protein-activity matrix of 3,000 regulatory proteins, where each cell has an inferred activity value for every protein in the matrix ^17, 20–22^.

Our study uniquely leverages computational protein expression inference for analysis of cell phenotypes in a cohort of CRC and further validates findings through multi-plex immunofluorescence (mIF) and traditional staining methods on a tissue microarray (TMA) composed of in-house samples from 50 EO- and 50-LOCRC patients (Figure 1A). Through this multi-modal approach, we demonstrate that the overall cellular composition of the TME is surprisingly similar between EO-CRC and LO-CRC, with the most notable difference being an increase in fibroblasts in EO-CRC, noted in a previous study ^23^. Amongst the epithelial compartment, we find a subpopulation of epithelial cells overrepresented in EO-CRC that are marked by increased expression of immune regulatory proteins TLR4, C5AR2, LILRB4, and CCR5. These findings suggest that EO-CRC may harbor a unique inflammatory epithelial compartment and enhanced stromagenesis.

**Figure 1:**
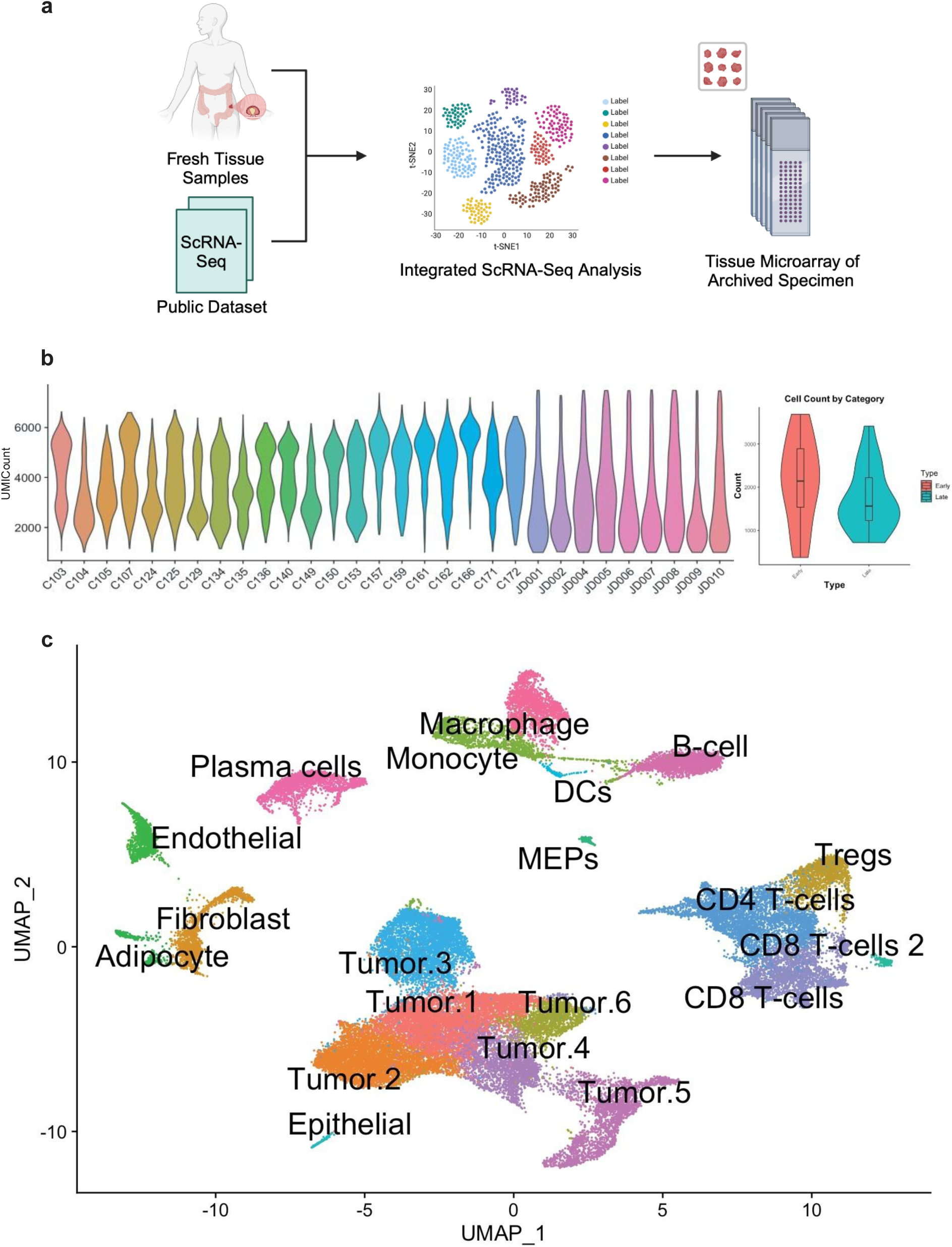
Experimental Workflow, Data Quality Metrics, and Single-Cell Transcriptomic Clustering. **a)** Experimental design. **b)** Distribution of Unique Molecular Identifiers (UMI) per cell, grouped by patient (left), distribution of post-quality control (QC) cell counts per patient, grouped by category, with no statistically significant difference in distribution. **c)** Uniform Manifold Approximation and Projection (UMAP) dimensionality reduction of entire combined ScRNA-seq dataset, with unsupervised clusters annotated by cell type.

## Results

### Patient Demographics and Characteristics

ScRNA-seq analysis was performed on an integrated cohort of 9 in-house patient samples and 21 patient samples obtained from a publicly available dataset that had not been previously analyzed by age ^24^. All tumors were taken from treatment-naive individuals with adenocarcinoma of the colon or rectum without known metastatic disease. All patients were considered Microsatellite Stable (MSS). Our in-house cohort consisted of 4 patients (44.4%) with EO-CRC (median age 43.5, range 42-46) and 5 patients (55.6%) with LO-CRC (median age 64, range 55-88). The public dataset included 7 EO-CRC patients (33.3%; median age 46, range 35-49) and 14 LO-CRC patients (66.7%; median age 68, range 57-90) who met inclusion criteria.

Demographic analysis across the combined cohort demonstrated a slight male predominance in the EO-CRC cohort (54.55 % male, 45.45% female; p=1.0), while LO-CRC demonstrated a marked male predominance (84.21% male, 15.79% female; p=0.012). While binomial test indicated significant overrepresentation of males in the LO-CRC cohort, there was no significant difference in gender distribution between the two groups by Fisher’s exact test (p=0.225). Anatomic distribution analysis confirmed previously reported trends, with EO-CR showing significant left-sided predominance (81.82% left-sided, 18.18% right-sided, p= 0.0386) compared to a more balanced distribution in LO-CRC (52.63% left-sided, 47.37% right sided; p = 1.0). There was no significant difference in tumor location distribution between the two groups by Fisher’s exact test (p= 0.140)

### Single-Cell Data Characteristics

ScRNA-seq yielded 42,931 cells cumulatively across patients after quality control filtering, with EO-CRC contributing 38.67% of the total cells, and no obvious outliers contributing exceedingly high or low cell count (Figure 1B). Quality metrics demonstrated robust data quality with 4000 median Unique Molecular Identifier (UMI) count per cell, and no significant batch effect by dataset of origin (Supplementary Figure S6). We initially clustered the data using Seurat R package on gene expression into distinct cell lineages. For each gene expression cluster, we used Algorithm for the Reconstruction of Accurate Cellular Networks (ARACNe) gene regulatory network inference algorithm to infer lineage-specific gene regulatory networks, as previously described ^17, 25, 26^. We then applied the VIPER protein activity algorithm to the initial gene expression data using the set of ARACNe regulatory networks, yielding a matrix of 2,748 regulatory proteins (genes annotated in the Gene Ontology (GO) molecular function database as transcription factors (TFs), co-TFs, or signaling molecules) across 42,931 cells. Unsupervised clustering on this matrix with resolution optimized by silhouette score yielded 20 distinct cellular populations (Figure 1C)^17^. Cell types for each cluster were labelled by majority-vote of single-cell level SingleR algorithm inferences ^27^. For validation studies, we utilized a tissue microarray (TMA) cohort of 100 additional patients, equally distributed between EO-CRC (n=50) and LO-CRC (n=50).

### Cellular Composition and Spatial Distribution of the Tumor Microenvironment

Comprehensive analysis of TME populations revealed both conserved and divergent features between EO-CRC and LO-CRC. While overall distribution of cells in the TME was similar between EO- and LO-CRC (Figure 2A), fibroblasts emerged as a significantly enriched population in EO-CRC, with mean cell frequency 3% vs 0.5% in LO-CRC, p=0.042 (Figure 2B). Detailed sub-clustering analysis identified Myofibroblast-like cancer-associated fibroblasts (myCAF) as the predominant fibroblast subtype (Figure 3A, 3B), comprising 92% and 93% of total fibroblasts in EO-CRC and LO-CRC respectively, with no difference in frequency of fibroblast sub-clusters in EO-CRC vs LO-CRC, suggesting conservation of fibroblast phenotypes despite differences in abundance (Figure 3C).

**Figure 2:**
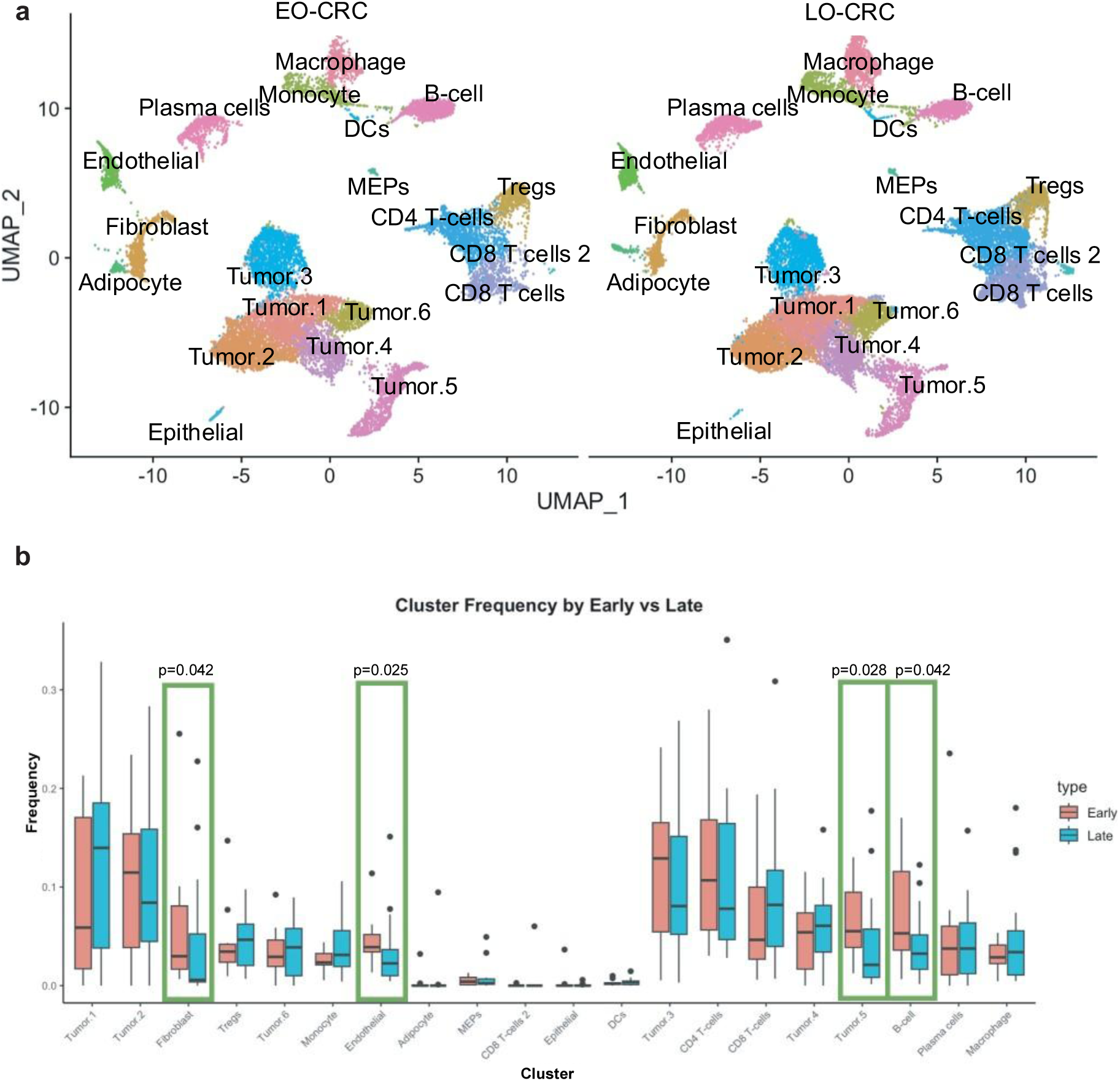
Comparison of Cellular Populations in EO-CRC and LO-CRC Identified by ScRNA-seq. **a)** UMAP dimensionality reduction of entire combined ScRNA-seq dataset, split by EO-CRC vs LO-CRC. **b)** Boxplot of cluster frequencies from Fig 1C grouped by patient type. Statistical significance of difference between EO-CRC and LO-CRC is assessed by Bonferroni-corrected Wilcox test, with statistically significant clusters outlined in green.

**Figure 3:**
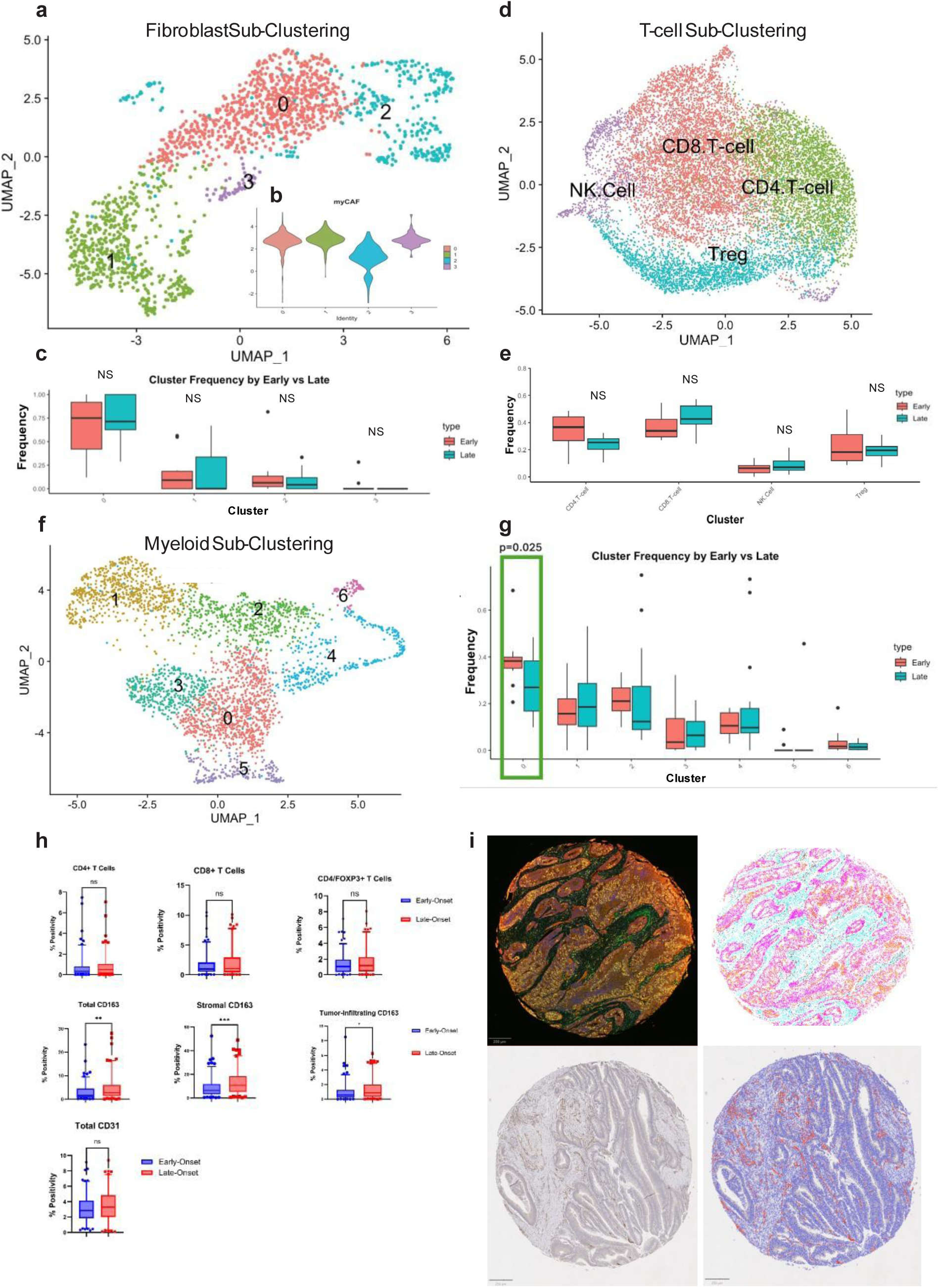
Cell Type-Specific Sub-Clustering and Tumor Microenvironment Composition in EO-CRC and LO-CRC. **a)** UMAP plot for sub-clustering of fibroblasts isolated from Figure 1C. **b)** Violin plot of GSEA Enrichment for myCAF vs iCAF gene signature, showing distribution across each fibroblast cell, grouped by cluster, where clusters 0, 1, and 3 are enriched for myCAF signature. **c)** Frequency plot comparing fibroblast sub-cluster frequencies by patient type. No statistically significant results. **d)** UMAP plot for sub-clustering of T-cells isolated from Figure 1C. **e)** Frequency plot comparing T-cell sub-cluster frequencies by EO- vs LO-CRC. No statistically significant results **f)** UMAP plot for sub-clustering of myeloid cells isolated from Figure 1C. **g)** Frequency plot comparing myeloid sub-cluster frequencies by EO- vs LO-CRC. Statistical significance of difference between EO-and LO-CRC is assessed by Bonferroni-corrected Wilcox test, with statistically significant clusters outlined in green. **h)** Quantification of TMA staining for immune and endothelial cell abundance in EO-and LO-CRC. **(Top row)** Quantification of CD4+ T cells, CD8+ T cells, and CD4+/FOXP3+ regulatory T cells (Tregs) revealed no significant differences between EO- and LO-CRC. **(Middle row)** CD163+ macrophages exhibited significant enrichment in LO-CRC: total CD163+ macrophages (p = 0.0029), stromal CD163+ macrophages (p = 0.0007), and tumor-infiltrating CD163+ macrophages (p = 0.0143). **(Bottom row)** CD31 staining for endothelial cells showed no significant difference between groups. Statistical significance was determined using a two-tailed Mann-Whitney U test. **i)** Representative Multiplex immunofluorescence (mIF) staining of TMA core (**top left**) with corresponding cell segmentation (**top right**). mIF staining was performed using Opal dyes: Opal 520 (CD8), Opal 540 (CD4), Opal 580 (Ki-67), Opal 620 (CD163), Opal 650 (FOXP3), and Opal 690 (EPCAM). Immunohistochemistry (IHC) staining for CD31 (**bottom left**) with corresponding cell segmentation (**bottom right**).

Immune cell profiling revealed subtle but significant variations between age groups. B cells showed modest enrichment in EO-CRC (5% vs 3% in LO-CRC, p=0.042), mirroring patterns observed in age-matched normal colon tissue (Figure 2B, Supplemental figure S5). This parallel suggests an age-related immune phenomenon rather than cancer-specific alterations. T cell populations, including CD4+, CD8+, and FOXP3+ regulatory T cells (Treg), showed remarkable consistency between groups across both ScRNA-seq and TMA validation cohorts (Figure 2B, 3D, 3E, 3H). Isolating lymphoid cells to sub-cluster further recapitulated CD4+, CD8+, Treg, and NK cell phenotypes but did not demonstrate any statistically significant difference in frequency between EO-CRC vs LO-CRC for these clusters, even as percent of lymphoid infiltrate (Figure 3D and 3E).

Myeloid cell analysis demonstrated age-specific compartmentalization. While ScRNA-seq globally showed no significant differences in frequency of myeloid cells between EO-CRC and LO-CRC, there was a general trend towards increased monocytes and macrophages in the LO-CRC samples (Figure 2B). Further monocytes sub-clustering revealed one sub-population of macrophages over-represented in EO-CRC as a fraction of total myeloid cells (Figure 3F, 3G). This sub-cluster exhibited an M2-like phenotype with activity of TREM2, APOE, and CSF1R expression (Supplemental Figure S2C, S2D), previously reported to associate with poorly prognostic immune-suppressive macrophage phenotype ^17^. Multiplex immunofluorescence (mIF) demonstrated elevated CD163+ macrophages in LO-CRC, particularly within stromal regions (10.57% vs 6.281% in EO-CRC, p=0.0007; Figure 3H, 3I). This stromal enrichment was complemented by a smaller but significant increase in tumor-adjacent regions (0.8905% vs 0.5688%, p=0.014; Figure 3H).

Endothelial cells presented an interesting methodological dichotomy. ScRNA-seq indicated significant enrichment in EO-CRC (p=0.025; Figure 2B), but subsequent IHC validation showed comparable frequencies (p=0.1509; Figure 3H), highlighting the importance of multi-modal validation approaches.

### Epithelial Compartment Reveals Unique Signature of EO-CRC

Unsupervised clustering of epithelial cells revealed six distinct transcriptional states preserved across age groups. Among these, the Tumor.5 cluster showed significant enrichment in EO-CRC (fold change=3.2, p=0.028; Figure 2B). This cluster was characterized in unsupervised analysis by elevated expression of immune-modulators, including TLR4, C5AR2, LILRB4, and CCR5 (Figure 4A), as well as by enrichment of interferon and TGF-beta signaling pathways (Figure 4B and 4C), and a number of druggable proteins including ERBB3, CEACAM6, and TOP2A (Supplemental Figure S3). Toll like receptor 4 (TLR4) is a transmembrane receptor that plays a central role in the innate immune response ^28^. Elevated expression of TLR4 has been associated with CRC as well as obesity-induced chronic inflammation ^29–32^. Further analysis of TLR4’s ARACNe-inferred regulatory targets revealed enrichment in complement signaling, epithelial-mesenchymal transition, and inflammatory response (Supplemental Figure S1). We further infer cell-cell receptor-ligand interactions involving the Tumor.5 cluster and other cell types. Interactions of interest for future validation experiments include engagement of LAG3 immune checkpoint on T-cells by LGALS3 on Tumor.5 cluster cells (Figure 5).

**Figure 4:**
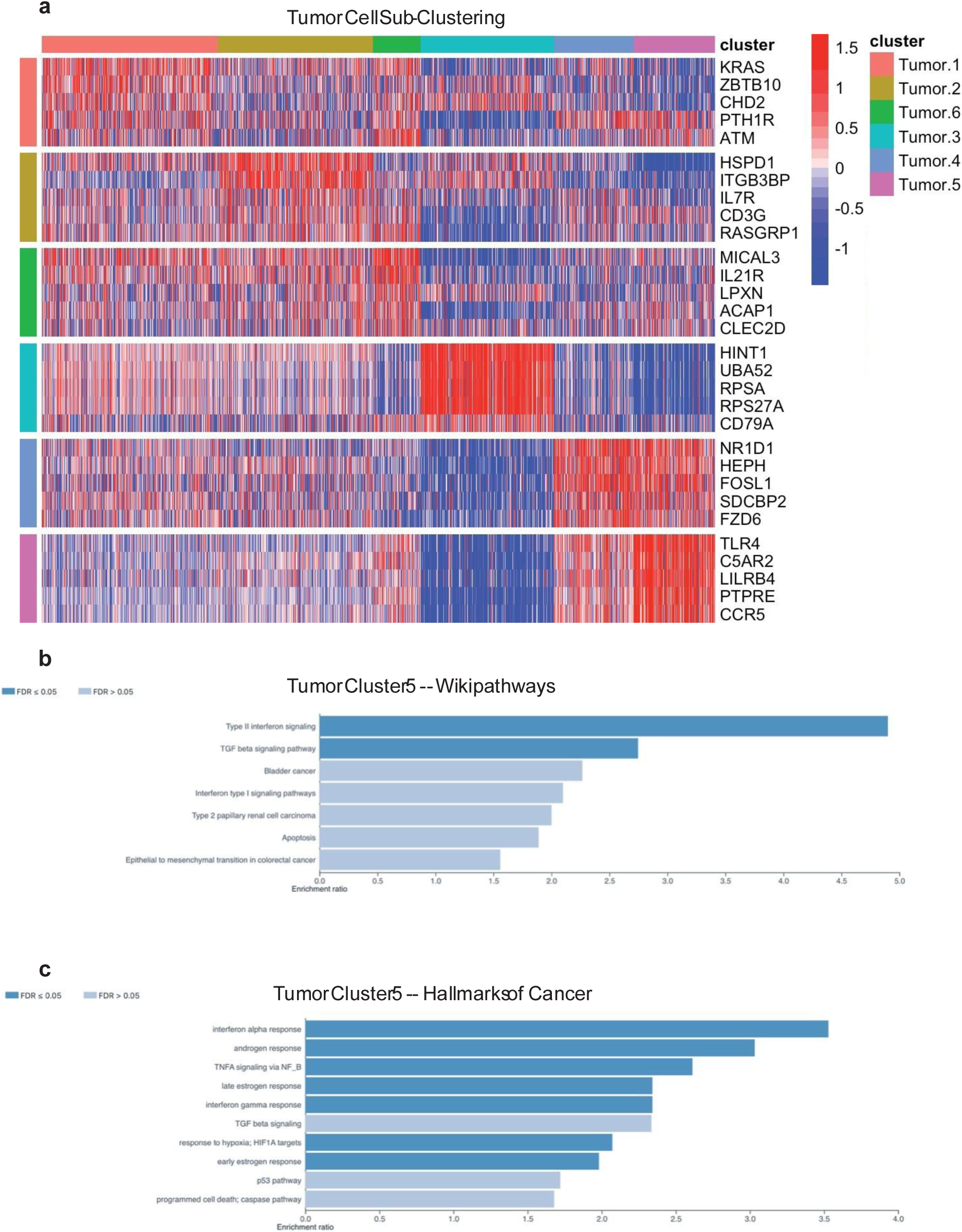
Defining Tumor Epithelial Cell Sub-Clusters. **a)** Heatmap of top 5 unsupervised proteins differentially upregulated by VIPER in tumor cell sub-clusters. **b)** Wikipathways pathway enrichment for differentially upregulated protein markers of Tumor.5 cell cluster, where statistically enriched pathways with FDR <0.05 are shown in dark blue. **c)** Hallmarks of Cancer pathway enrichment for differentially upregulated protein markers of Tumor.5 cell cluster, where statistically enriched pathways with FDR <0.05 are shown in dark blue.

**Figure 5:**
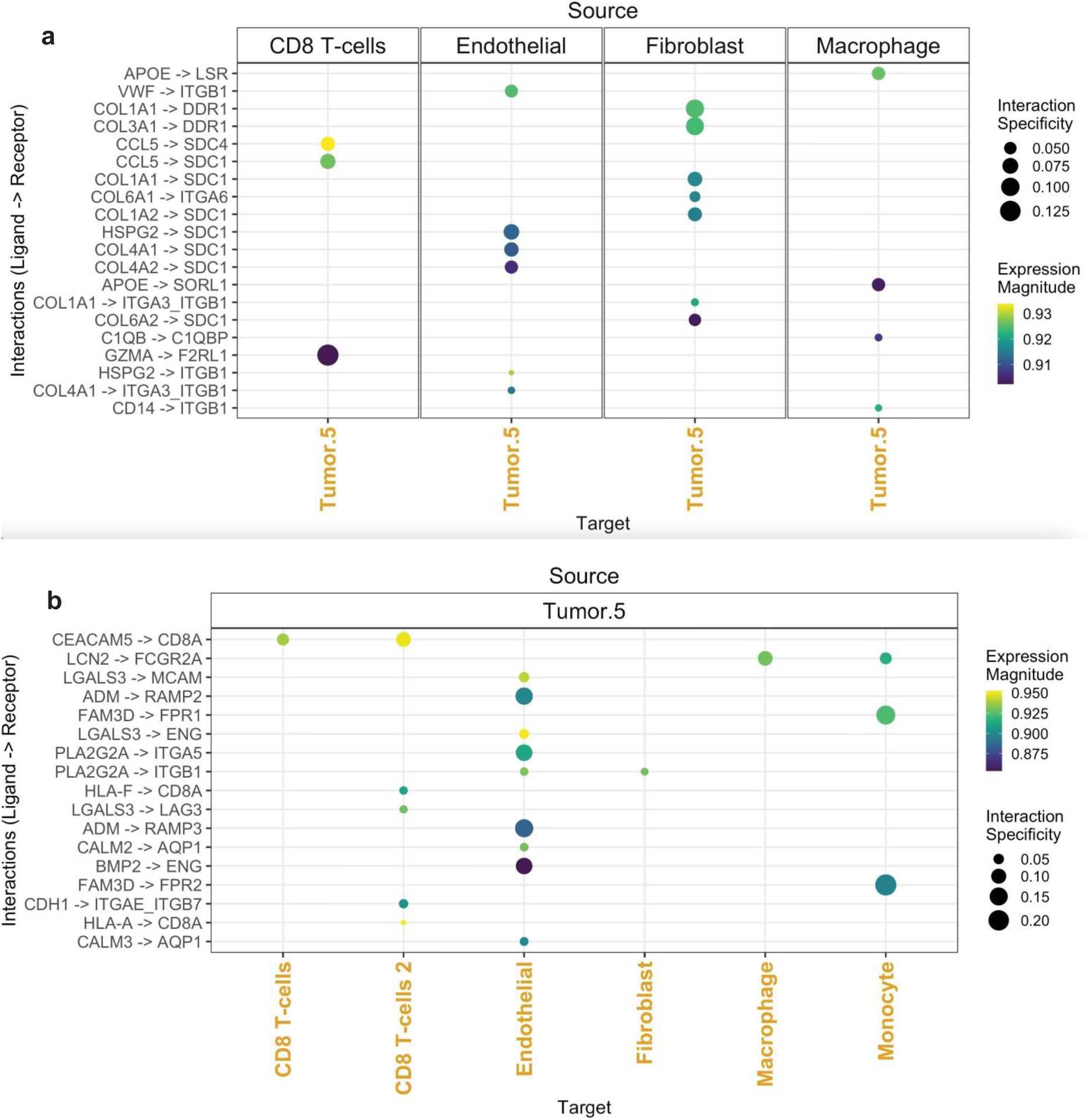
Inference of Tumor.5 Cell Cluster Receptor-Ligand Signaling. **a)** Inference by LIANA of receptor-ligand interactions involving Tumor.5 cell cluster as the receptor-expressing cell cluster, showing all interactions with statistical significance <0.05. **b)** Inference by LIANA of receptor-ligand interactions involving Tumor.5 cell cluster as the ligand-expressing cell cluster, showing all interactions with statistical significance <0.05.

## Discussion

The rising incidence of sporadic (non-hereditary) EO-CRC over the past several decades has driven recent research to elucidate its molecular underpinnings. Despite advances in genomic and transcriptional profiling, consistent molecular distinctions between EO-CRC and LO-CRC have remained elusive. Through detailed single-cell transcriptional profiling and inferred protein expression analysis, we have identified several important findings, including similar tumor microenvironment and an increase in a distinct subpopulation of epithelial cells in EO-CRC characterized by enhanced expression of inflammatory pathways associated with diet and the microbiome.

Contrary to previous reports suggesting an “immune-cold” phenotype in EO-CRC, our analysis of the TME revealed remarkable similarities between EO-CRC and LO-CRC immune landscapes ^15, 33^. The comparable densities of CD4+, CD8+, and FOXP3+ T cells between age groups suggest that these tumors are immunologically similar. Interestingly, our findings of increased B cells in EO-CRC align with those from a recent single-cell study of EO-CRC in an East Asian cohort^16^. However, our study provides additional context by demonstrating that B-cell enrichment is not unique to EO-CRC but is also present in normal colon tissue from younger, healthy individuals (Supplemental Figure S5). This suggests that age-associated immune variation, rather than a tumor-specific phenomenon, may underlie this observation, highlighting the importance of considering baseline immune differences when interpreting TME alterations in EO-CRC. Our analysis revealed some variations in stromal components, most notably an increase in cancer-associated fibroblasts (CAFs) in EO-CRC and modest differences in CD163+ macrophage distribution. The increased CAF presence in EO-CRC is consistent with previous reports but showed no specific subtype enrichment in myCAF or inflammatory cancer-associated fibroblasts (iCAFs), suggesting a broader stromal response rather than a unique fibroblast program in EO-CRC ^23^.

When investigating the epithelial compartment, we find an increase in epithelial Tumor.5 cluster, marked by several inflammatory genes associated with colorectal cancer, most notably TLR4. The sub-stratification of tumor cells necessary to make this finding would not have been possible with raw gene expression analysis and emerged with the aid of VIPER protein activity inference, a unique contribution of this manuscript. TLR4 serves as a critical mediator of both microbiota-driven inflammation and dietary-induced metabolic stress ^32, 34^. Certain saturated fatty acids have been found to bind to TLR4 and activate TLR4-mediated inflammatory pathways in a similar manner to bacterial lipopolysaccharides (LPS) ^34^. This dual responsiveness to both dietary factors and microbial signals positions TLR4 as a potential integrator of environmental influences in EO-CRC pathogenesis. Supporting this hypothesis, studies have shown that high-fat diets rich in palmitic acid increase TLR4 expression and enhance CRC proliferation in murine models ^32^. This TLR4-dependent mechanism is independent of changes to tumor infiltrating immune cells, consistent with our observation of conserved immune cell profiles between EO- and LO-CRC ^32^.

TLR4’s role in colorectal carcinogenesis has been well documented through multiple studies ^29, 35, 36^. In the context of inflammatory bowel disease (IBD)-associated CRC, Fukata et al. demonstrated that TLR4 is overexpressed in human tumor tissue. Additionally, TLR4-knockout (KO) mice showed significant protection against tumor development in a murine model of colitis-associated neoplasia ^35^. Subsequent studies revealed that TLR4 signaling promotes carcinogenesis through increased epithelial reactive oxygen species (ROS) production in a microbiota-dependent manner ^36^. This mechanism was elegantly demonstrated using Villin-TLR4 mice that overexpress TLR4 throughout the gastrointestinal tract and develop tumors significantly faster than Wild-Type (WT) mice. Notably, this accelerated tumor development was prevented in germ-free conditions, and WT mice receiving fecal transplants from Villin-TLR4 mice showed increased tumor burden compared to those receiving WT fecal material ^36^. These findings suggest that upregulation of TLR4 and intestinal dysbiosis are both necessary to form this positive-feedback loop that results in CRC development and progression.

In addition to TLR4, further analysis of the Tumor.5 cluster reveals increased expression of several genes associated with tumor progression and microenvironment remodeling. This cluster exhibits high expression of C-C chemokine receptor type 5 (CCR5), a chemokine receptor implicated in fibroblast recruitment and tumor-associated inflammation^37, 38^. Notably, CCR5 signaling has been linked to poor prognosis in CRC, where it promotes an immunosuppressive microenvironment and facilitates CAF activation^39^. The upregulation of CCR5, alongside inflammatory mediators such as C5AR2, LILRB4, and CXCL10, suggests that this epithelial subpopulation may play a critical role in shaping the EO-CRC microenvironment through immune-modulatory and stromal interactions.

Our findings provide new evidence for a possible cell-intrinsic mechanism with enhanced stromagenesis present in EO-CRC. Several limitations of our study, however, warrant discussion and point to important future directions. While our single-cell analysis revealed a distinct epithelial population in EO-CRC marked by TLR4 expression and other inflammatory mediators, our study design was cross-sectional, limiting our ability to establish causality between dietary factors, TLR4 expression, and tumor development. Future longitudinal studies tracking dietary patterns, microbiome composition, and inflammatory mediator expression in young adults could help establish the temporal relationship between these factors and EO-CRC risk. Additionally, while our patient cohort was carefully selected to include only mismatch repair (MMR)-proficient, non-metastatic tumors, the sample size is modest. Larger multi-center cohorts will be needed to validate our findings across diverse patient populations. Furthermore, although we identified increased TLR4 and other inflammatory marker expression in EO-CRC tumor cells, we did not directly assess the underlying mechanisms involved. Future studies using patient-derived organoids and murine models could help elucidate the direct effects of TLR4 signaling on tumor cell behavior. Ultimately, these limitations provide a roadmap for future studies to advance targeted prevention strategies and therapeutic interventions for EO-CRC.

## Methods

### Single Cell RNA Sequencing Tissue Processing

In-house clinical samples were obtained from human colorectal cancer resection specimens under (IRB AAAT8778). Upon receipt of tissue, specimens were cut into 1cm x 1cm sections and viably frozen in 90% FBS + 10% DMSO. On the day of ScRNA-seq, tissue was flash thawed and minced into small pieces using a scalpel. Tissue was washed x3 using Primocin (InvivoGen, ant-pm) and then digested using Miltenyi Tumor Dissociation kit per manufacturer instructions (Miltenyi Biotec, 130-095-929). Digest was then filtered through a 70-micron strainer and washed x2 to ensure removal of the digestion buffer. Cells were counted using Countess 3 (Invitrogen) and brought on ice to the Single Cell RNA-Sequencing Core at Columbia University where 10x protocol was performed per manufacturer’s instructions.

Briefly, samples were processed for single-cell gene expression capture using the 10X Chromium 3’ Library and Gel Bead Kit (10x Genomics), following the manufacturer’s user guide at the Columbia University Single Cell Analysis Core. After GelBead in-Emulsion reverse transcription (GEM-RT) reaction, 12–15 cycles of polymerase chain reaction (PCR) amplification were performed to obtain cDNAs used for RNAseq library generation. Libraries were prepared following the manufacturer’s user guide and sequenced on Illumina NovaSeq 6000 Sequencing System or the Illumina NovaSeqX System. Single-cell RNASeq data were processed with Cell Ranger software at the Columbia University Single Cell Analysis Core. Illumina base call files were converted to FASTQ files with the command “cellranger mkfastq.” Cell Ranger performed default filtering for quality control, and produced for each sample a barcodes.tsv, genes.tsv, and matrix.mts file containing counts of transcripts for each sample, such that expression of each gene is in terms of the number of unique molecular identifiers (UMIs) tagged to cDNA molecules corresponding to that gene.

### Single-Cell RNA-seq Public Datasets

Publicly available ScRNA-seq data was obtained from the Gene Expression Omnibus (GEO) under accession number GSE178341, originally published by Pelka et al., 2021 (PMID: 34450029). This dataset was used for integration with in-house samples to compare EO-CRC with LO-CRC. This dataset was also used to compare adjacent normal tissue between Young (<50 years) and old (≥50 years) patient cohorts. Publicly available ScRNA-seq data of healthy colon tissue was obtained from the Human Cell Atlas (HCA) Data Portal under the project “A Single-Cell Atlas of the Normal and Malignant Human Colon” [https://explore.data.humancellatlas.org/projects/fde199d2-a841-4ed1-aa65-b9e0af8969b1]

### Single-Cell RNA-seq Gene Expression Processing

Gene Expression UMI count matrices for each sample were processed in R using the Seurat SCTransform command to perform a regularized negative binomial regression based on the 3000 most variable genes. Samples were then combined and batch-corrected using the FindIntegrationAnchors and IntegrateData functions in Seurat v4, with default parameters [https://pubmed.ncbi.nlm.nih.gov/31178118/]. The combined dataset was then clustered by Resolution-Optimized Louvain Clustering Algorithm described in [https://pubmed.ncbi.nlm.nih.gov/34019793/], such that cluster resolution was selected to maximize average silhouette score across bootstrapped sub-samples of the full dataset. Within each cluster metaCells were computed for downstream regulatory network inference by summing SCTransform-corrected template counts for the 10 nearest neighbors of each cell by Pearson correlation distance. The resulting dataset was projected into its first 50 principal components using RunPCA function in Seurat, and further reduced into a 2-dimensional visualization space using the RunUMAP function with method umap-learn and Pearson correlation as the distance metric between cells. Differential Gene Expression between clusters was computed by the MAST hurdle model for single-cell gene expression modeling [https://genomebiology.biomedcentral.com/articles/10.1186/s13059-015-0844-5], as implemented in the Seurat FindAllMarkers command, with log fold change threshold of 0.5 and minimum fractional expression threshold of 0.25, indicating that the resulting gene markers for each cluster are restricted to those with log fold change greater than 0 and non-zero expression in at least 25% of the cells in the cluster.

### Regulatory Network Inference

From each gene expression cluster, metaCells were computed by summing SCTransform-corrected template counts for the 10 nearest neighbors of each cell by Pearson correlation distance. 200 metaCells per cluster were sampled to compute a regulatory network from each cluster in each patient. All regulatory networks were reverse engineered by the ARACNe algorithm. ARACNe was run with 100 bootstrap iterations using 1785 transcription factors (genes annotated in gene ontology molecular function database as GO:0003700, “transcription factor activity,” or as GO:0003677, “DNA binding” and GO:0030528, “transcription regulator activity,” or as GO:0003677 and GO:0045449, “regulation of transcription”), 668 transcriptional cofactors (a manually curated list, not overlapping with the transcription factor list, built upon genes annotated as GO:0003712, “transcription cofactor activity,” or GO:0030528 or GO:0045449), 3455 signaling pathway related genes (annotated in GO biological process database as GO:0007165, “signal transduction” and in GO cellular component database as GO:0005622, “intracellular” or GO:0005886, “plasma membrane”), and 3620 surface markers (annotated as GO:0005886 or as GO:0009986, “cell surface”). ARACNe is only run on these gene sets so as to limit protein activity inference to proteins with biologically meaningful downstream regulatory targets, and we do not apply ARACNe to infer regulatory networks for proteins with no known signaling or transcriptional activity for which protein activity may be difficult to biologically interpret. Parameters were set to zero DPI (Data Processing Inequality) tolerance and MI (Mutual Information) p value threshold of 10^-8, computed by permuting the original dataset as a null model.

### Protein Activity Inference and Re-Clustering

Protein activity was inferred for all cells by running the VIPER algorithm with the full set of ARACNe networks across on the SCTransform-scaled and Anchor-Integrated gene expression signature of single cells. Because the SCTransform-scaled gene expression signature is already normalized, VIPER normalization parameter was set to “none.” This resulted in a matrix of 2,748 proteins with successfully inferred activity across all cells. Activity for each protein in each cell is inferred as a Normalized Enrichment Score (NES). The VIPER-Inferred Protein Activity matrix was loaded into a Seurat Object with CreateSeuratObject, then projected into its first 50 principal components using the RunPCA function and further reduced into a 2-dimensional visualization space using the RunUMAP function with method umap-learn and Pearson correlation as the distance metric between cells. Differential Protein Activity between unsupervised clusters identified by resolution-optimized Louvain was computed using T-test with Benjamini-Hochberg multiple-testing correction, and top proteins for each cluster were ranked by fold-change (Figure 4A, S2A-C, S4). Pathway Enrichment of differentially active protein sets was assessed by WebGestalt (https://academic.oup.com/nar/article/52/W1/W415/7684598) (Figure 4B-C), and activity of druggable proteins was assessed by overlap with the set of all proteins with known drug modulator in DrugBank (https://pubmed.ncbi.nlm.nih.gov/37953279/), subsetting to drugs where VIPER protein normalized enrichment score corresponds to corrected p-value < 0.01 on average in at least one tumor cell cluster (Supplemental Figure S3).

### Semi-Supervised Cell Type Calling and Sub-Clustering

For each single cell gene expression sample, cell-by-cell identification of cell types was performed using the SingleR package (https://www.nature.com/articles/s41590-018-0276-y) and the preloaded Blueprint-ENCODE reference, which includes normalized expression values for 259 bulk RNASeq samples generated by Blueprint and ENCODE from 43 distinct cell types representing pure populations of stroma and immune cells (https://pubmed.ncbi.nlm.nih.gov/22955616/). The SingleR algorithm calculates correlation between each individual cell and each of the 259 reference samples and then assigns both a label of the cell type with highest average correlation to the individual cell and a p value computed by Wilcox test of correlation to that cell type compared to all other cell types. Cell labels are filtered to those with p<0.05, and intersected with unsupervised clustering labels, such that localization of SingleR labels is highly concordant with the protein activity unsupervised clustering. Unsupervised Protein Activity Clusters determined by the resolution-optimized Louvain algorithm are labeled as a particular cell type based on the most-represented SingleR cell type label within that cluster.

For cell clusters identified as fibroblast, myeloid, and lymphoid, sub-clustering was performed on isolated protein activity matrix, by recomputing RunPCA and RunUMAP and re-computing unsupervised resolution-optimized Louvain clusters. For T-cell sub-clusters, SingleR cell type labels remained strongly concordant with unsupervised cluster groups (Figure 3D). For T-cell and myeloid cell sub-clusters, phenotyping was augmented by visualizing inferred activity of manually selected gene panels as well (supplemental figure S2D,E). For fibroblast sub-clusters, we additionally performed cell-by-cell Gene Set Enrichment Analysis (GSEA) of previously reported iCAF and myCAF gene markers, first reported in pancreatic cancer context and publicly available at (https://pmc.ncbi.nlm.nih.gov/articles/PMC6727976/) – visualization of gene set enrichment in violin plot grouped by cluster identified one sub-cluster as predominantly iCAF-like and the rest as myCAF-like (Figure 3B). Receptor-Ligand interactions between cell types (Figure 5) were inferred using LIANA algorithm (https://pubmed.ncbi.nlm.nih.gov/35680885/).

### Tissue Microarray (TMA) Preparation

This work was performed in the Molecular Pathology Shared Resource of the Herbert Irving Comprehensive Cancer Center at Columbia University. Formalin-fixed, paraffin-embedded (FFPE) tumor samples were used to construct a TMA for immunohistochemical and multiplex immunofluorescence analysis. Representative tumor regions were identified by a board-certified pathologist and cored using a 2mm punch from donor blocks. Tissue cores were precisely arrayed into a recipient paraffin block (Beecher Instruments), ensuring uniform spatial distribution. The completed TMA block was sectioned at 5 µm, mounted on charged glass slides, and stored at room temperature until staining.

### Immunohistochemistry (IHC) Staining and Analysis

Tissue microarray (TMA) sections were subjected to immunohistochemical (IHC) staining to assess protein expression of CD31. Formalin-fixed, paraffin-embedded (FFPE) TMA sections were deparaffinized and rehydrated through graded ethanol washes. Antigen retrieval was performed using a citrate buffer (pH 6.0) in a water bath at 95°C for 15 minutes with an additional 15 minutes at room air. Slides were blocked using 3% hydrogen peroxide, Avidin, Biotin, and Starting Block T20 for 15 minutes each.

Sections were incubated with primary antibody anti-CD31 (Clone C31/1395R, 1ug/ml, Cat# 5175-RBM11-P0, NeoBiotechnologies) diluted in 1% BSA/0.3% triton X in PBX at room temperature for 2 hours followed by incubation with biotinylated Goat Anti-Rabbit antibody (Vectra, VL_BA-1000, 1:200 dilution) at 4°C for 30 minutes. Slides were developed using ABC amplification complex for 30 minutes at 37°C followed by DAB (3,3’-diaminobenzidine) substrate for 10 minutes. Slides were transferred to H20 to stop the reaction and were counterstained with hematoxylin.

Stained slides were scanned at high resolution using Leica Aperio AT2 whole slide digital scanner at 40x, and quantification was performed using Qupath (0.5.1-x64). Automated cell segmentation was validated by manual review. Each core was independently evaluated by two blinded reviewers. Cores with poor staining quality or insufficient tissue were excluded from the analysis. Statistical analysis of cell counts was conducted using a two-tailed Mann-Whitney U test.

### Multiplex Tissue Imaging and Spatial Analysis

The Leica Bond RX Fully Automounted Research Stainer performed the heating, dewaxing, and staining functions for all TMA slides. Heating and dewaxing functions were necessary prior to staining under the approved Leica 7-Color Double Dispensing protocol to ensure proper adhesion of the tissue onto the glass slide. Under the staining protocol, tissues were treated with six primary antibodies and 10X Spectral DAPI (Akoya, Cat #FP1490). Along with the primary antibody, tissues were subsequently treated with an HRP-conjugated secondary anti-mouse antibody (Akoya, ARH1001EA) and an opal fluorophore.

Akoya 1X Antibody Diluent Block (Akoya, Cat# ARD1001EA) was used to dilute antibodies to desired concentrations while opals were resuspended in DMSO before further dilutions with 1X Plus Automation Amplification Diluent. The following antibody-opal order and pairings were used to stain and image early and late onset TMA slides: anti-FoxP3 Antibody (Abcam Cat # ab20034, Clone 236A/E7, 1/200) and Opal 650 (Akoya, Cat # FP1496001KT, 1/175), anti-CD4 Antibody (Abcam, Cat # ab133616, Clone EPR6855, 1/1000) and Opal 540 (Akoya, Cat # FP1494001KT, 1/200), anti-CD163 Antibody (Leica, Cat # NCL-L-CD163, Clone 10D6, 1/400) and Opal 620 (Akoya, Cat # FP1495001KT, 1/200), anti-CD8 Antibody (Leica, Cat # NCL-L-CD8-4B11-L, Clone 4B11, 1/50) and Opal 520 (Akoya, Cat # FP1487001KT, 1/200), anti-Ki67 Antibody (Abcam, Cat # AB15580, polyclonal, 1/200) and Opal 570 (Akoya, Cat # FP1488001KT, 1/175), anti-EPCAM Antibody (Abcam, Cat #ab223582, Clone EPR20532-225, 1/500) and Opal 690 (Akoya, Cat # FP1497001KT, 1/200).

Multispectral imaging was performed on the Vectra Polaris PhenoImager at 20X magnification for each TMA slide following cover slipping with No. 1 Superslip Micro Glasses (EMS Cat #72236-50) and Antifade Mounting Medium (Vectashiled Cat #H-1000).

Analysis was performed utilizing the inForm 3.0 software. Multispectral fields underwent unmixing to differentiate overlapping signals, autofluorescence, and background as opals often express in multiple filters when adjacent opal channels are utilized. Tissue segmentation was performed to highlight areas exhibiting elevated levels of EPCAM, low levels of EPCAM, and areas of non-interest. Individual cells were segmented with aid from nuclear markers DAPI and FoxP3 as well as membrane markers CD4, EPCAM, Ki67, and CD8 to further aid in nuclear splitting. The typical intensity for each marker along with the splitting sensitivity, and minimum nuclear size were set for both nuclear and assisting components. Cellular phenotyping for each of the six markers was also performed as was phenotyping for cells of non-interest and the colocalized expression of Ki67/EPCAM, FoxP3/CD4, Ki67/CD4, Ki67/CD8. After the inForm algorithm had been trained to perform unmixing, tissue segmentation, cell segmentation, and cell phenotyping, individual TMAs were analyzed through a batch analysis. Data from each TMA was later merged to highlight the number of cellular phenotypes identified per tissue segmented region.

### Quantification and Statistical Analysis

Quantitative and statistical analyses for ScRNA-seq were performed using the R computational environment and packages described above. Differential gene expression was assessed at the single-cell level by the MAST single-cell statistical framework as implemented in Seurat v4, and differential VIPER protein activity was assessed by T-test, each with Benjamini-Hochberg multiple-testing correction. Comparisons of cell frequencies were performed by non-parametric Wilcox rank-sum test. In all cases, statistical significance was defined as an adjusted p value less than 0.05. Details of statistical tests used can be found in the corresponding figure legends.

Quantitative and statistical analyses of TMA staining were performed using GraphPad Prism (Version 10.4.1). Comparisons between EO-CRC and LO-CRC groups were assessed using the Mann-Whitney U test for non-parametric distributions. Comparison of patient demographics, including gender distribution and tumor sidedness between EO-CRC and LO-CRC cohorts, were performed using two-tailed Fisher’s exact test and binomial test as appropriate. Statistical significance was defined as p value less than 0.05%.

## Acknowledgements

This work was supported by the Columbia Cancer Training Program for Resident-Investigators, NIH grant # 1R38CA277850-01A1. This work was supported by several Core Facilities at Columbia University, including, the Human Immune Monitoring Core, Molecular Pathology Shared Resource of the Herbert Irving Comprehensive Cancer Center, and the Single Cell Core at Columbia Genome Center, supported by NIH grant #P30CA013696. The Single Cell Core at Columbia Genome Center is further supported by the NIH grant #UL1TR001873.

## Author Contributions

D.A.C. led the study design, data analysis, and manuscript development, coordinating input from all co-authors to finalize submission. A.O. completed single-cell RNA-sequencing (ScRNA-Seq) analysis, provided input on computational approaches, assisted with figure and manuscript generation. L.J., Y.M., S.L., F.C. assisted with bioinformatics analysis. Y.P., B.D., S.V., CW.C., contributed to methodology development and analysis. T.L., D.D., CN.B., A.S., C.L., N.L., J.H., contributed to data interpretation and provided critical feedback on the manuscript. K.D., C.R., V.J., assisted with immunohistochemistry (IHC) and multiplex immunofluorescence (mIF) experiments. E.B. assisted with patient recruitment and tissue procurement. J.G. supervised the study, provided conceptual guidance, and edited the manuscript. All authors read and approved the final manuscript.

## Materials & Correspondence

Correspondence and material requests should be directed to Dr. Joel Gabre at. For access to data, materials, or additional information regarding this study, please contact Dr. Joel Gabre at the provided email. Requests will be considered in accordance with institutional policies and data-sharing guidelines.

## References

1. Bhandari, A., Woodhouse, M. & Gupta, S. Colorectal cancer is a leading cause of cancer incidence and mortality among adults younger than 50 years in the USA: a SEER-based analysis with comparison to other young-onset cancers. J. Investig. Med. 65, 311–315 (2017).

2. Vuik, F. E. R. et al. Increasing incidence of colorectal cancer in young adults in Europe over the last 25 years. Gut 68, 1820–1826 (2019).

3. Siegel, R. L., Wagle, N. S., Cercek, A., Smith, R. A. & Jemal, A. Colorectal cancer statistics, 2023. CA Cancer J. Clin. 73, 233–254 (2023).

4. Mauri, G. et al. Early-onset colorectal cancer in young individuals. Mol. Oncol. 13, 109–131 (2019).

5. Bailey, C. E. et al. Increasing Disparities in the Age-Related Incidences of Colon and Rectal Cancers in the United States, 1975-2010. JAMA Surg. 150, 17–22 (2015).

6. Anderson, J. C. & Samadder, N. J. To screen or not to screen adults 45-49 years of age: That is the question. Am. J. Gastroenterol. 113, 1750–1753 (2018).

7. Ullah, F., Pillai, A. B., Omar, N., Dima, D. & Harichand, S. Early-Onset Colorectal Cancer: Current Insights. Cancers (Basel) 15, 3202 (2023).

8. Stoffel, E. M. et al. Germline Genetic Features of Young Individuals with Colorectal Cancer. Gastroenterology 154, 897 (2018).

9. Siegel, R. L. et al. Colorectal Cancer Incidence Patterns in the United States, 1974-2013. J. Natl. Cancer Inst. 109, (2017).

10. Akimoto, N. et al. Rising incidence of early-onset colorectal cancer — a call to action. Nat. Rev. Clin. Oncol. 18, 230–243 (2021).

11. Patel, S. G., Karlitz, J. J., Yen, T., Lieu, C. H. & Boland, C. R. The rising tide of early-onset colorectal cancer: a comprehensive review of epidemiology, clinical features, biology, risk factors, prevention, and early detection. Lancet Gastroenterol. Hepatol. 7, 262–274 (2022).

12. O’Sullivan, D. E. et al. Risk Factors for Early-Onset Colorectal Cancer: A Systematic Review and Meta-analysis. Clin. Gastroenterol. Hepatol. 20, 1229–1240.e5 (2022).

13. Cohen, D., Rogers, C., Gabre, J. & Dionigi, B. The Young: Early-Onset Colon Cancer. Clin. Colon Rectal Surg. (2024) doi:10.1055/S-0044-1787883.

14. Tang, J. et al. Molecular characteristics of early-onset compared with late-onset colorectal cancer: a case controlled study. Int. J. Surg. 110, 4559–4570 (2024).

15. Ugai, T. et al. Immune cell profiles in the tumor microenvironment of early-onset, intermediate-onset, and later-onset colorectal cancer. Cancer Immunol. Immunother. 71, 933–942 (2022).

16. Li, G.-M. et al. Single-cell RNA sequencing reveals heterogeneity in the tumor microenvironment between young-onset and old-onset colorectal cancer. Biomolecules 12, 1860 (2022).

17. Obradovic, A. et al. Single-cell protein activity analysis identifies recurrence-associated renal tumor macrophages. Cell 184, 2988–3005.e16 (2021).

18. Obradovic, A. et al. Immunostimulatory cancer-associated fibroblast subpopulations can predict immunotherapy response in head and neck cancer. Clin. Cancer Res. 28, 2094–2109 (2022).

19. Mundi, P. S. et al. A transcriptome-based precision oncology platform for patient-therapy alignment in a diverse set of treatment-resistant malignancies. Cancer Discov. 13, 1386–1407 (2023).

20. Vlahos, L. et al. Systematic, protein activity-based characterization of single cell state. bioRxiv 2021.05.20.445002 (2021) doi:10.1101/2021.05.20.445002.

21. Wang, X., He, Y., Zhang, Q., Ren, X. & Zhang, Z. Direct comparative analyses of 10X Genomics Chromium and Smart-seq2. Genomics Proteomics Bioinformatics 19, 253–266 (2021).

22. Alaqeeli, O. A comparison of dropout rate of three commonly used single cell RNA-sequencing protocols. Biotechnol. Biotechnol. Equip. 38, (2024).

23. Furuhashi, S. et al. Spatial profiling of cancer-associated fibroblasts of sporadic early onset colon cancer microenvironment. *NPJ Precis*. Oncol. 7, 1–19 (2023).

24. Pelka, K. et al. Spatially organized multicellular immune hubs in human colorectal cancer. Cell 184, 4734–4752.e20 (2021).

25. Margolin, A. A. et al. ARACNE: an algorithm for the reconstruction of gene regulatory networks in a mammalian cellular context. BMC Bioinformatics 7 Suppl 1, S7 (2006).

26. Hao, Y. et al. Integrated analysis of multimodal single-cell data. Cell 184, 3573–3587.e29 (2021).

27. Aran, D. et al. Reference-based analysis of lung single-cell sequencing reveals a transitional profibrotic macrophage. Nat. Immunol. 20, 163–172 (2019).

28. Kim, H.-J., Kim, H., Lee, J.-H. & Hwangbo, C. Toll-like receptor 4 (TLR4): new insight immune and aging. Immun. Ageing 20, 67 (2023).

29. Cammarota, R. et al. The tumor microenvironment of colorectal cancer: stromal TLR-4 expression as a potential prognostic marker. J. Transl. Med. 8, (2010).

30. Semlali, A. et al. Expression and polymorphism of toll-like receptor 4 and effect on NF-κB mediated inflammation in colon cancer patients. PLoS One 11, e0146333 (2016).

31. Kim, F. et al. Toll-like receptor-4 mediates vascular inflammation and insulin resistance in diet-induced obesity. Circ. Res. 100, 1589–1596 (2007).

32. Hu, X. et al. Toll-like receptor 4 is a master regulator for colorectal cancer growth under high-fat diet by programming cancer metabolism. Cell Death Dis. 12, 1–13 (2021).

33. Andric, F. et al. Immune Microenvironment in Sporadic Early-Onset versus Average-Onset Colorectal Cancer. Cancers (Basel) 15, (2023).

34. Rogero, M. M. & Calder, P. C. Obesity, inflammation, toll-like receptor 4 and fatty acids. Nutrients 10, 432 (2018).

35. Fukata, M. et al. Toll-like receptor-4 promotes the development of colitis-associated colorectal tumors. Gastroenterology 133, 1869–1881 (2007).

36. Burgueño, J. F. et al. Epithelial TLR4 signaling activates DUOX2 to induce Microbiota-driven tumorigenesis. Gastroenterology 160, 797–808.e6 (2021).

37. Sasaki, S., Baba, T., Shinagawa, K., Matsushima, K. & Mukaida, N. Crucial involvement of the CCL3-CCR5 axis-mediated fibroblast accumulation in colitis-associated carcinogenesis in mice: CCL3-CCR5 axis in colon carcinogenesis. Int. J. Cancer 135, 1297–1306 (2014).

38. Tanabe, Y., Sasaki, S., Mukaida, N. & Baba, T. Blockade of the chemokine receptor, CCR5, reduces the growth of orthotopically injected colon cancer cells via limiting cancer-associated fibroblast accumulation. Oncotarget 7, 48335–48345 (2016).

39. Suarez-Carmona, M. et al. CCR5 status and metastatic progression in colorectal cancer. Oncoimmunology 8, e1626193 (2019).

