## Supplemental Figures S1 - S6 for "Single-Cell RNA Sequencing and Inferred Protein Activity Analysis Reveal a Distinct Tumor Phenotype in Early-Onset Colorectal Cancer Patients"

### Supplemental Figure S1

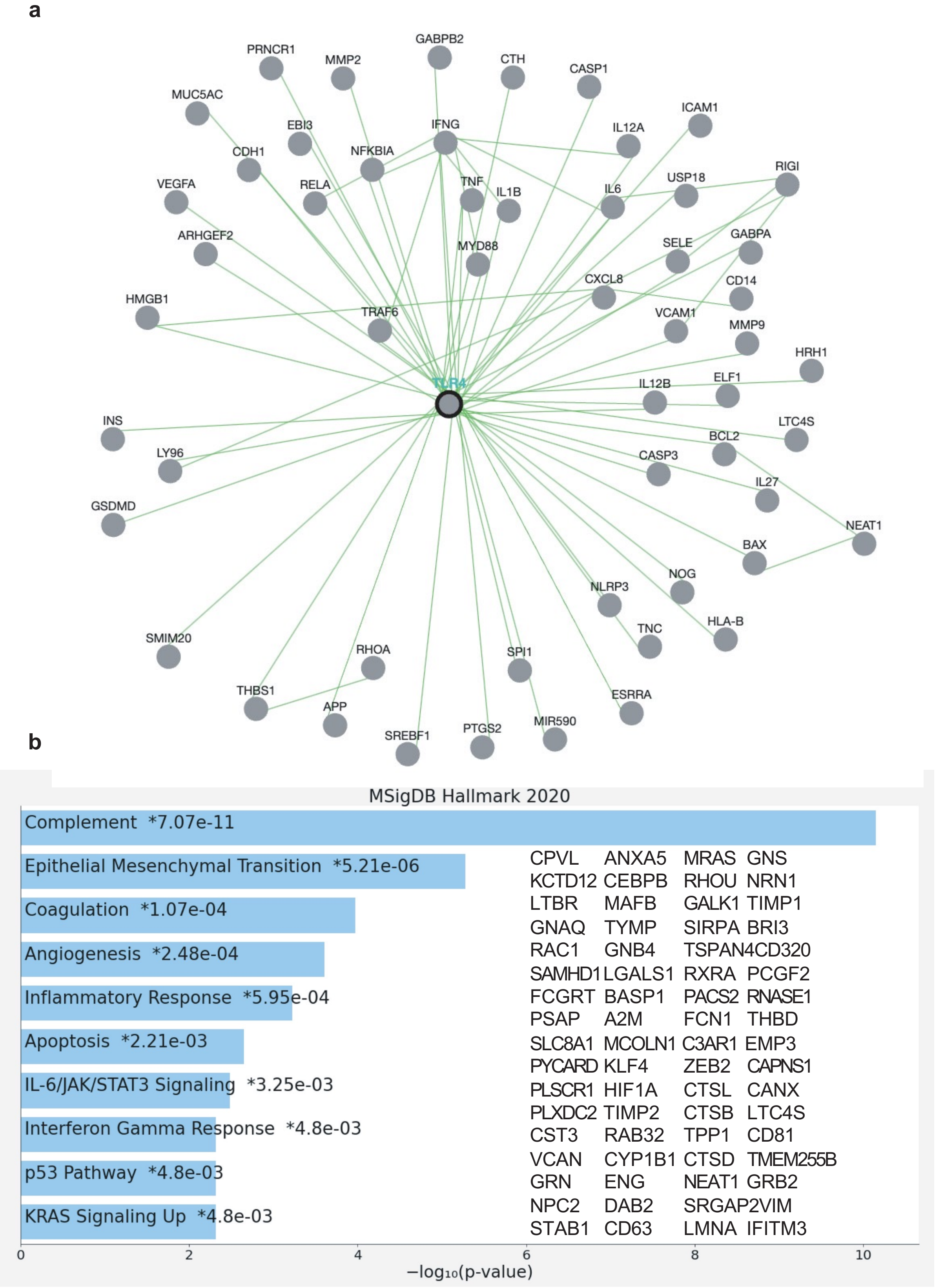

**Supplemental Figure S1: Regulatory Network and Functional pathway Analysis of TLR4 Target Genes**

**a)** Network graph of downstream target genes regulated by TLR4. **b)** Hallmarks of Cancer Pathway Enrichment for downstream target genes regulated by TLR4. Inset text with comprehensive gene list of inferred downstream targets upregulated by TLR4.

### Supplemental Figure S2

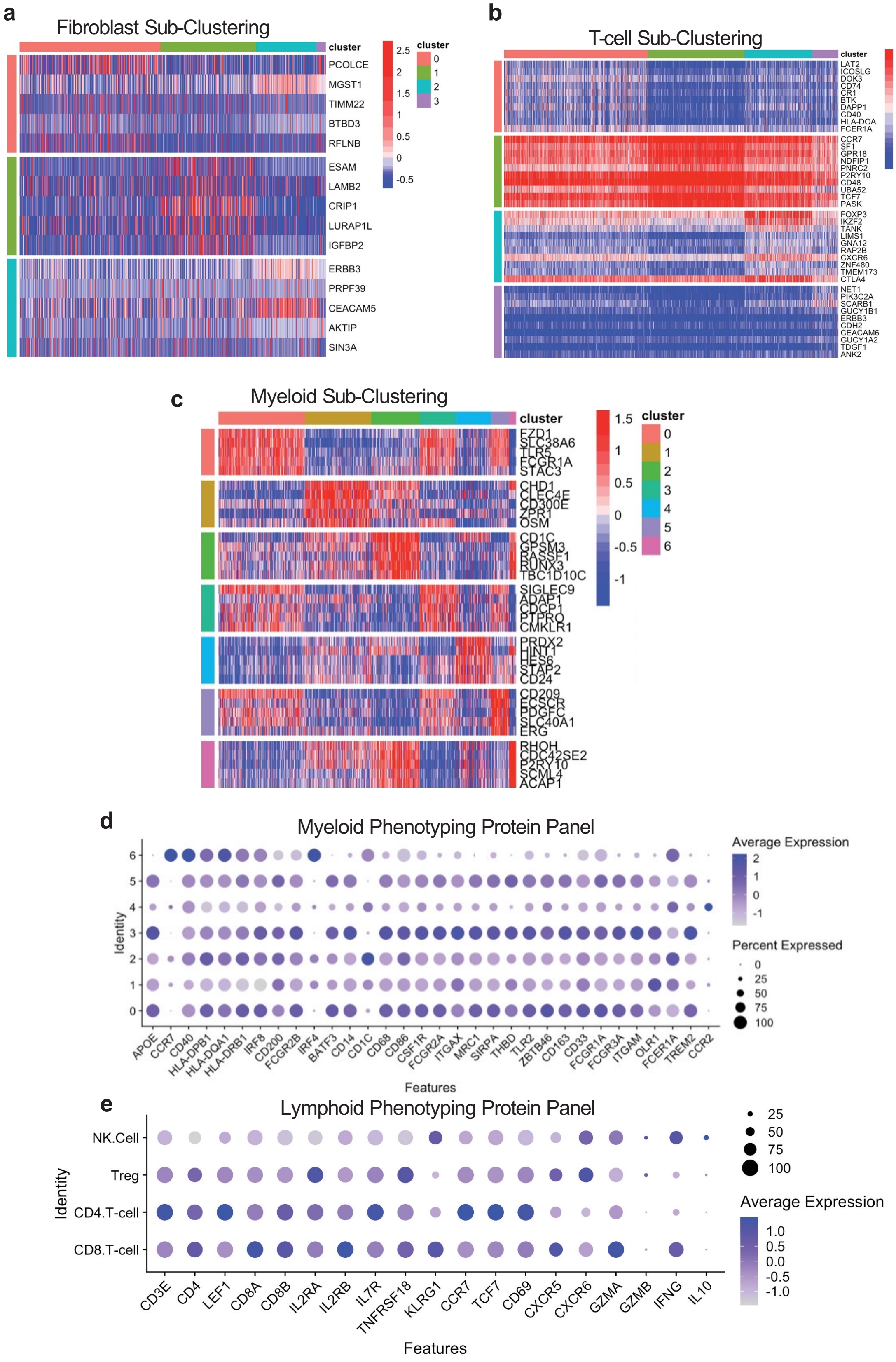


7

**Supplemental Figure S2: VIPER-Inferred Protein Activity Across Fibroblast, T-Cell, and Myeloid Sub-Clusters**

**a)** Heatmap of top 5 unsupervised proteins most differentially upregulated by VIPER in each Fibroblast sub-cluster corresponding to Figure 3A. **b)** Heatmap of top10 unsupervised proteins most differentially upregulated by VIPER in each T-cell sub-cluster corresponding to Figure 3D **c)** Heatmap of top5 unsupervised proteins most differentially upregulated by VIPER in each Myeloid sub-cluster corresponding to Figure 3F. **d)** Dot plot showing inferred protein activity of manually-selected genes across each Myeloid sub-cluster from C. **e)** Dot plot showing inferred protein activity of manually-selected genes across each T-cell sub-cluster from B.

Supplemental Figure S3

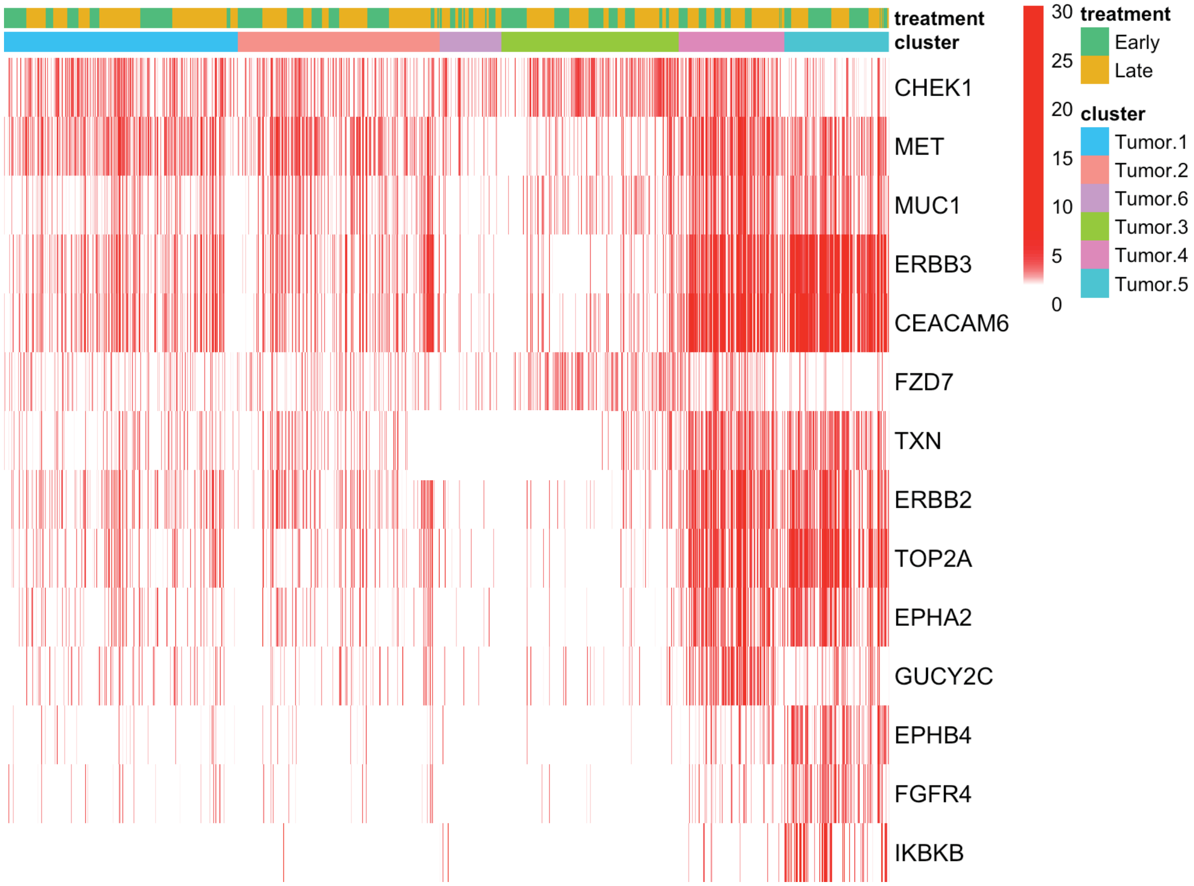


**Supplemental Figure S3: Enrichment of Druggable Targets Across Tumor Cell Sub-Clusters**

Heatmap of -log10(p-values) for enrichment of druggable proteins upregulated with mean statistical significance $<0.01$ in at least one tumor cell sub-cluster.

### Supplemental Figure S4

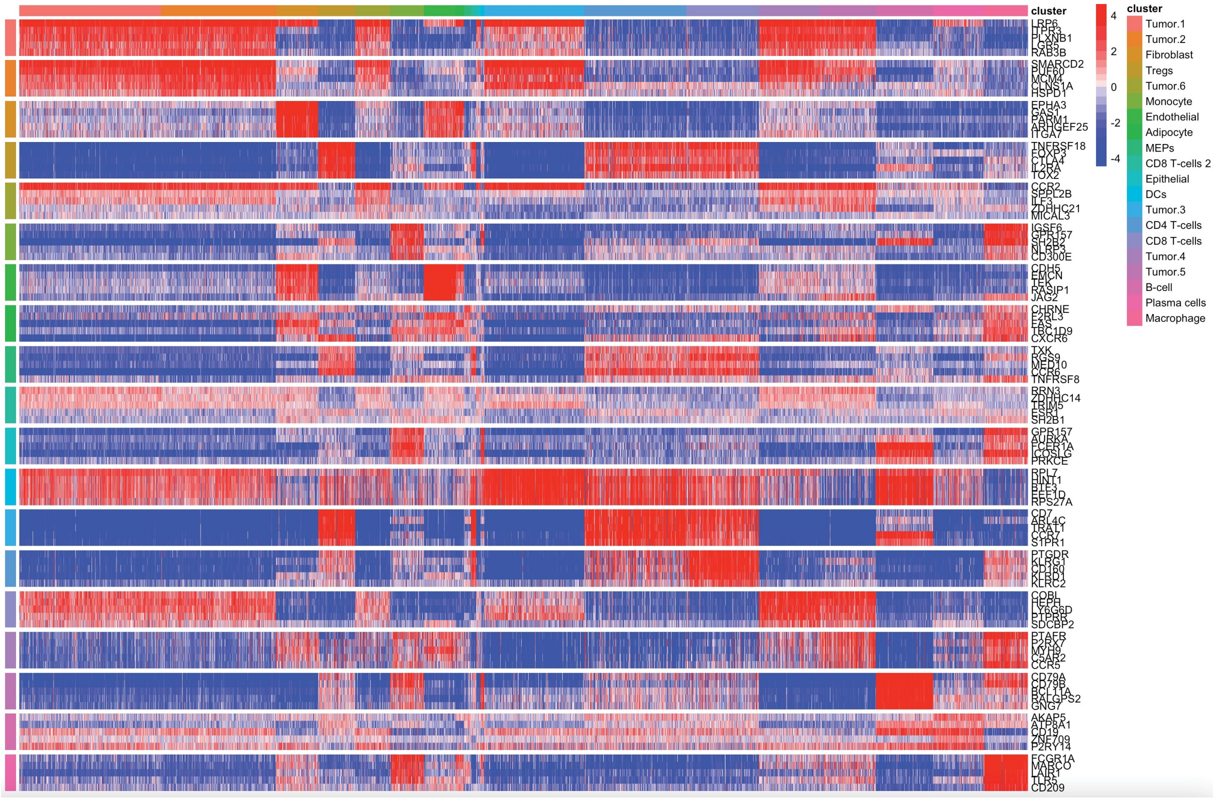


**Supplemental Figure S4: VIPER-Inferred Differentially Upregulated Proteins Across Single-Cell clusters**

Heatmap of top 5 unsupervised proteins differentially upregulated by VIPER in each cluster from Figure 1C.

### Supplemental Figure S5

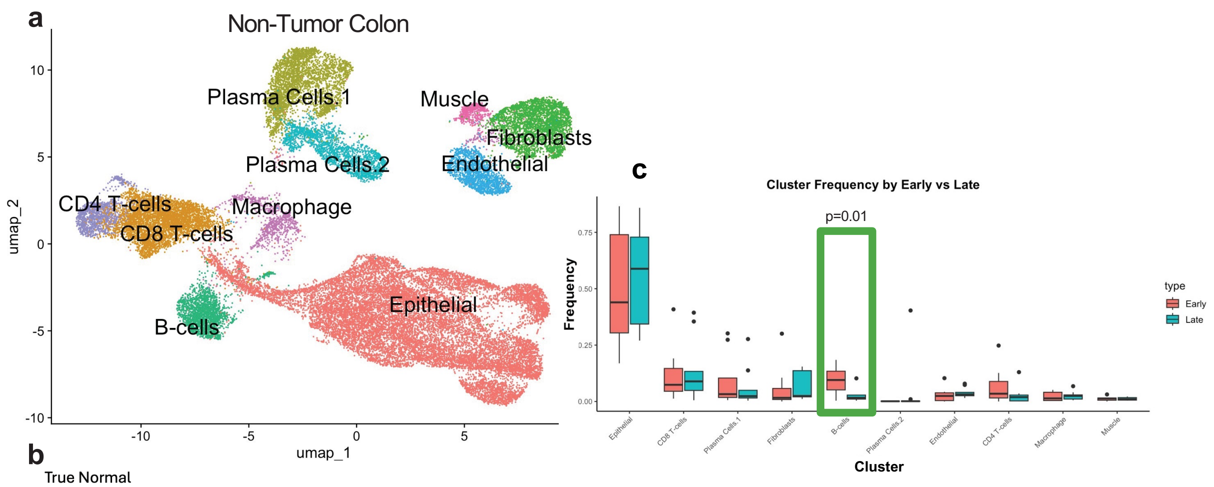


b

True Normal

| Sample | A26 | A30 | A32 | A33 | A34 | A38 | A39 |
|---|---|---|---|---|---|---|---|
| Cell Count | 1333 | 2511 | 4581 | 3704 | 4823 | 2253 | 2922 |
| Type | Late | Early | Early | Early | Late | Late | Early |

Non-Tumor

| Sample | C107 | C124 | C125 | C129 | C134 | C135 | C136 | C140 | C157 | C162 |
|---|---|---|---|---|---|---|---|---|---|---|
| Cell Count | 907 | 2299 | 1617 | 823 | 871 | 472 | 798 | 839 | 1698 | 3363 |
| Type | Late | Late | Early | Early | Late | Late | Late | Early | Late | Early |

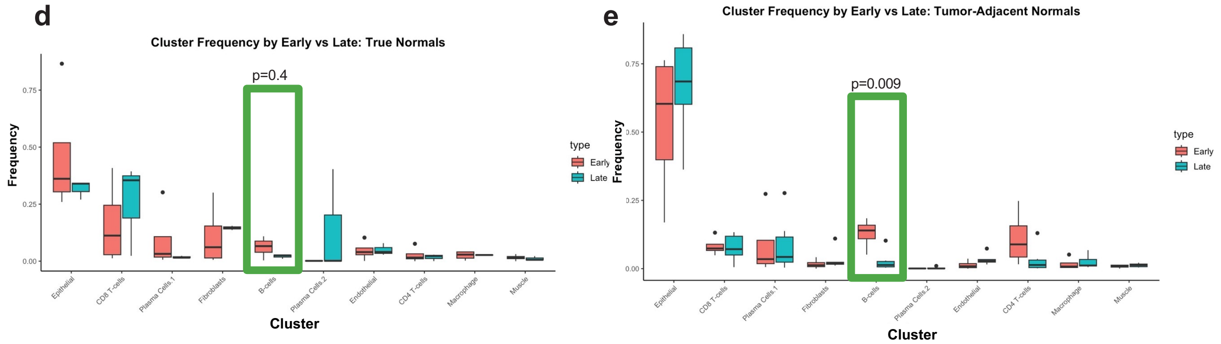


**Supplemental Figure S5: Single-Cell Characterization of Non-Tumor Colon Tissue Across Early and Late-Onset Samples**

**a)** UMAP clustering for inferred protein activity of integrated non-tumor colon scRNA-Sequencing data, with cell types of unsupervised clusters annotated by singleR as in Figure 2A. **b)** Table of number of cells post-QC filtering for each sample included in A, annotated as tumor-adjacent normal tissue vs true normal and as early vs late (based on same age criteria as EO-CRC and LO-CRC). **c)** Frequency plot showing differential abundance of each cluster in early vs late normal tissue, in aggregate of all samples. p-values assessed by Wilcox test with Benjamini-Hochberg multiple-testing correction. Only statistically significant difference found for B-cells. **d)** Frequency plot showing differential abundance of each cluster in early vs late normal tissue, as in C, subset to only true normal samples. Due to low sample size, no statistically significant differences were observed. **e)** Frequency plot showing differential abundance of each cluster in early vs late normal tissue, as in C, subset to tumor-adjacent normal samples. p-values assessed by Wilcox test with Benjamini-Hochberg multiple-testing correction. Only statistically significant difference found for B-cells.

**Supplemental Figure S6**

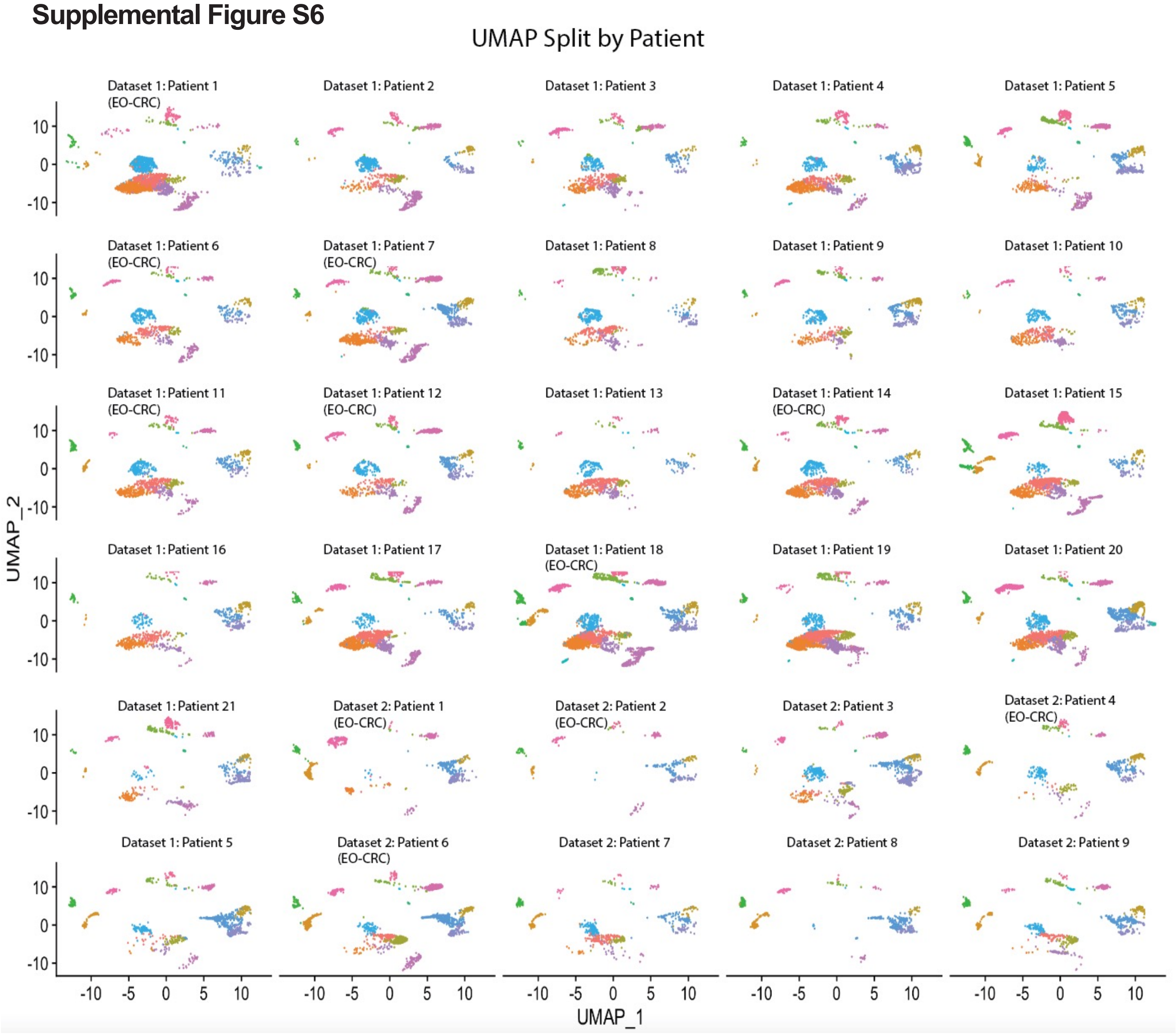


**Supplemental Figure S6: Assessment of Batch Effects in Unsupervised Clustering Across Individual Patients.**

UMAP plot of combined unsupervised clustering, as in Figure 1C, split by individual patient, showing lack of significant batch effect / patient-specific clusters.
